# Identification of pre-existing microbiome and metabolic vulnerabilities to escalation of oxycodone self-administration and identification of a causal role of short-chain fatty acids in addiction-like behaviors

**DOI:** 10.1101/2022.07.23.501268

**Authors:** Sierra Simpson, Adam Kimbrough, Gregory Peters, Emma Wellmeyer, Rio Mclellan, Natalie Walker, Haoyu Jia, Sharon Hu, Mohini Iyer, Varshini Sathish, Sharona Sedighim, Marsida Kallupi, Molly Brennan, Lisa Maturin, Talyn Hughes, Tristin Xie, Veronika Espinoza, Lieselot Carrette, Lauren C. Smith, Jonathan Seaman, Leah C. Solberg Woods, Abraham A. Palmer, Giordano DeGuglielmo, Olivier George

**Affiliations:** Department of Psychiatry, University of California San Diego; Department of Basic Medical Sciences, College of Veterinary Medicine, Purdue University; Department of Psychology, University of Texas El Paso; National Center for Complementary and Integrative Health, National Institute of Health; Department of Internal Medicine, Section on Molecular Medicine, Wake Forest University School of Medicine; Institute for Genomic Medicine, University of California San Diego

## Abstract

The gut brain axis is thought to play a role in behavior and physiological responses through chemical, immunological, and metabolite signaling. Antibiotics, diet, and drugs can alter the transit time of gut contents as well as the makeup of the microbiome. Heterogeneity in genetics and environment are also well-known factors involved in the initiation and perpetuation of substance use disorders. Few viable genetic or biological markers are available to identify individuals who are at risk of escalating opioid intake. Primarily, the addiction field has focused on the nervous system, limiting the discovery of peripheral factors that contribute to addiction. To address this gap, we characterized the microbiome before and after drug exposure, and after antibiotics depletion in male and female heterogenous stock rats to determine if microbiome constituents are protective of escalation. We hypothesized that individuals that are prone to escalation of opioid self-administration will have distinct microbial and metabolic profiles. The fecal microbiome and behavioral responses were measured over several weeks of oxycodone self-administration and after antibiotic treatment. Antibiotic treatment reduces circulating short-chain fatty acids (SCFA) by depleting microbes that ferment fiber into these essential signaling molecules for the gut-brain axis. Depletion of the microbiome increased oxycodone self-administration in a subpopulation of animals (Responders). Supplementation of SCFAs in antibiotic depleted animals decreased elevated oxycodone self-administration. Phylogenetic functional analysis reveals distinct metabolic differences in the subpopulations of animals that are sensitive to antibiotic depletion and animals rescued by SCFA supplementation. In conclusion, this study identifies pre-existing microbiome and metabolic vulnerabilities to escalation of oxycodone self-administration, demonstrates that escalation of oxycodone self-administration dysregulates the microbiome and metabolic landscape, and identifies a causal role of short-chain fatty acids in addiction-like behaviors.

## Introduction

Opioids are an important pain management tool, however, opioid misuse and opioid use disorder represent a major health problem in the United States with a significant increase (15-30%) in overdose deaths over the past two years (Volkow and Blanco, 2021; NIDA, 2021). Recent evidence indicates that interactions between the gut microbiome and central nervous system are bidirectional and may contribute to addiction-related disorders (Cryan et al., 2019; Patterson et al., 2014; Simpson et al., 2021a; Simpson et al., 2020b; Simpson et al., 2021b). A side effect of opioids is disruption of gut motility through activation of peripheral opioid receptors that in turn alter the resident microbiota. In the clinic, patients are often prescribed opioids in conjunction with antibiotics. The mechanism of action of antibiotics can exaggerate the deleterious effects of opioids on the gut and deleteriously shift the microbiome when patients are most susceptible to the escalation of opioid use (Clegg et al., 2021; Freedman et al., 2021). However, the effect of the interaction of opioid and antibiotic use on the microbiome and metabolic pathways is poorly known, and it is unclear which microbiome and metabolic vulnerabilities may contribute to escalation of oxycodone self-administration. Indeed, human studies are very challenging due to the difficulty in enrolling patients prior opioid misuse, long-term monitoring of opioid use, the use of qualitative instead of qualitative measures, the presence of bias in self reporting, and the heterogeneity of patients with opioid use disorder. To address this issue, we leveraged a powerful translational model using heterogeneous stock rats with state-of-the art characterization of oxycodone self-administration and investigated the microbiome and metabolic landscape before and after escalation of oxycodone self-administration as well as after antibiotic treatments and short-chain fatty acid supplementation. We hypothesized that the microbiome contributes to pre-existing addiction vulnerability and that the disruption of the microbiome with antibiotics may further increase the vulnerability to addiction-like behaviors, including the escalation of oxycodone self-administration, and that analysis of the microbiome may lead to discovery of novel biomarkers relevant to predicting individual addiction liability.

To explore the intersection of the microbiome and opioid addiction-like behavior, the present study used male and female heterogenous stock (HS) rats, which exhibit high genetic diversity, obtained from the Oxycodone Biobank (Carrette et al., 2021). HS rats were created from combinatorial breeding of eight inbred strains followed by a colony maintenance protocol to minimizes inbreeding. After 70+ generations, the resulting animals’ chromosomes are a mosaic of the founders’ haplotypes, representing a powerful tool for translational studies (Woods and Mott, 2017). Animals were initially subset by intake (High versus Low) after 21-day of long access (12 h/day) to oxycodone self-administration to determine if microbiome composition prior to drug or antibiotic exposure was associated with distinct microbial profiles or functions and to explore which microbes/functions were altered following exposure to oxycodone. The metabolic profile of the microbes was analyzed throughout the experiment (PICRUST2 (Douglas et al., 2020), MetaCyC (Caspi et al., 2016)). Then the microbiome of subset of the animals was depleted by non-absorbable antibiotics to determine if depletion of SCFA producers would increase intake. A subgroup of animals that were sensitive to antibiotic depletion (ABX Responders) were treated with a SCFA cocktail which ameliorated the increased intake observed following antibiotic depletion in a subpopulation of animals (SFCA Responders). While repletion studies have been carried out in other drugs of abuse, this is the first time that repletion of SCFA in antibiotic depleted animals has been carried out in freely behaving individuals capable of choosing to take the substance. In addition, animals represented both sexes in a strain of rat that mimics the variability of drug taking behavior observed in the human population.

## Materials and Methods

### Methods

Drug naïve animals were implanted with intravenous catheters into the jugular vein at 8 weeks of age. The animals were allowed to recover for one week before operant self-administration training began. 1 week of short access training (2h) of oxycodone self-administration was allowed before moving to long access sessions (12h). The animals performed 21 days of long access self-administration before depletion of the microbiome was initiated. Animals were divided to match intake across three groups 1) Water (no abx, no SCFA treatment), 2) ABX (abx treatment for the duration of the experiment, 3) ABX+SCFA (abx treatment, then abx + SCFA treatment). At the start of the ABX treatment, both ABX and ABX+SCFA groups were depleted using a cocktail of non-absorbable antibiotics in their drinking water, the water group were given regular drinking water for the duration of the experiment. Following 16 days of antibiotic depletion, animals that received ABX were then divided evenly by intake and maintained on antibiotics, with half of the animals receiving additional supplementation of SCFAs. Prolonged administration of oxycodone leads to tolerance and physical dependence as shown by withdrawal symptoms upon the cessation of the drug. Withdrawal scores and the von Frey pain threshold test were taken at the time of withdrawal. Tolerance was measured by tail immersion at 55 C. At the end of the study rats were sacrificed and perfused 24h into withdrawal.

### Animals

Adult male (*n* = 34) and female (*n* = 34) Heterogenous stock (HS) rats were housed in groups of two with a 12:12 hour light/dark cycle, light on at 6:00 pm and off at 6:00 am in a humidity-controlled vivarium with *ad libitum* access to tap water and food pellets (PJ Noyes Company, Lancaster, NH, USA). The HS rat colony (N:NIH-HS) was first established by the NIH in 1984 using the following inbred strains: ACI/N,BN/SSN, BUF/N, F344/N, M520/N, MR/N, WKY/N, WN/N (Hansen and Spuhler, 1984). All procedures were conducted in strict adherence to the National Institutes of Health *Guide for the Care and Use of Laboratory Animals* and were approved by the Institutional Animal Care and Use Committee of Scripps Research. At the time of testing, subjects’ body weight ranged between 280-350 for females, and 350-400 g for males.

### Drugs

Oxycodone (Sigma Aldrich, St. Louis, MO) was dissolved in 0.9% sodium chloride (Hospira, Lake Forest, IL, USA) and administered via intravenous catheter in both short access and long access sessions at a dose of (150 μg/0.1 ml/kg). Antibiotic doses were given at the following concentrations in drinking water - Bacitracin (0.5 mg/ml), Neomycin (2 mg/ml), Vancomycin (0.2 mg/ml) and Pimaricin (1.2 μg/ml) similar to previous studies (Kiraly et al., 2016a) These agents were specifically selected because they characterized as being non-absorbed from the intestine and have been previously reported to only have efficacy in depleting the microbiome when administered orally (Simpson et al., 2020b; Kiraly et al., 2016a). This cocktail allows for maximal reduction of gut bacteria and minimal off-target effects. The antibiotic mixture was changed every two days and rats were weighed weekly to ensure body weight was maintained. Short chain fatty acids were given at the following concentrations – acetate (67.5 mM), butyrate (40 mM), and propionate (25.9 mM).

### Progressive Ratio

At the end of the ShA and LgA phases, a progressive ratio (PR) schedule of reinforcement was used to assess the break point in lever-press responding, a measure of the reinforcing value of a reward (Stafford et al., 1998; Hodos, 1961). The PR response requirements necessary to receive a single drug dose increased according to this equation: [5e ^(injection numbers x 0.2)^] – 5, resulting in the following progression of response requirements: 1, 2, 4, 6, 9, 12, 15, 20, 25, 32, 40, 50, 62, 77, 95, 118, 145, 178, 219, 268, etc. The breakpoint was defined as the last ratio attained by the rat prior to a 60 min period during which a ratio was not completed. The performance under progressive ratio was repeated two times after the LgA phase.

### Fecal Collection and 16S rRNA gene sequencing

At times marked in Figure 1A (BSL, LGA, ABX), rats were held by the base of the tail so that the front paws were on a solid surface and the back paws were lifted off the surface. The back paws were gently moved up and down off the surface until a fresh fecal pellet was released. The pellet was collected immediately into a sterile tube, put on dry ice, then frozen at -80°C for storage until sequencing. The pellets were then sent for extraction and sequencing by the UCSD Microbiome Sequencing Core. At the time of DNA extraction, feces were thawed and extracted with the Qiagen DNeasy powersoil kit. The samples are then amplified by PCR in triplicate and then pooled. The 16S V4 gene was amplified using universal primers (525F-806R). Each sample was normalized to 240 ng per sample and purified. After purification, the A260/A280 ratio of the final pool was recorded to ensure purity, with a tolerance range of 1.8-2.0. The barcoded amplicons from all of the samples were normalized, pooled to construct the sequencing library, and sequenced using an Illumina MiSeq sequencer.

**Figure 1.**
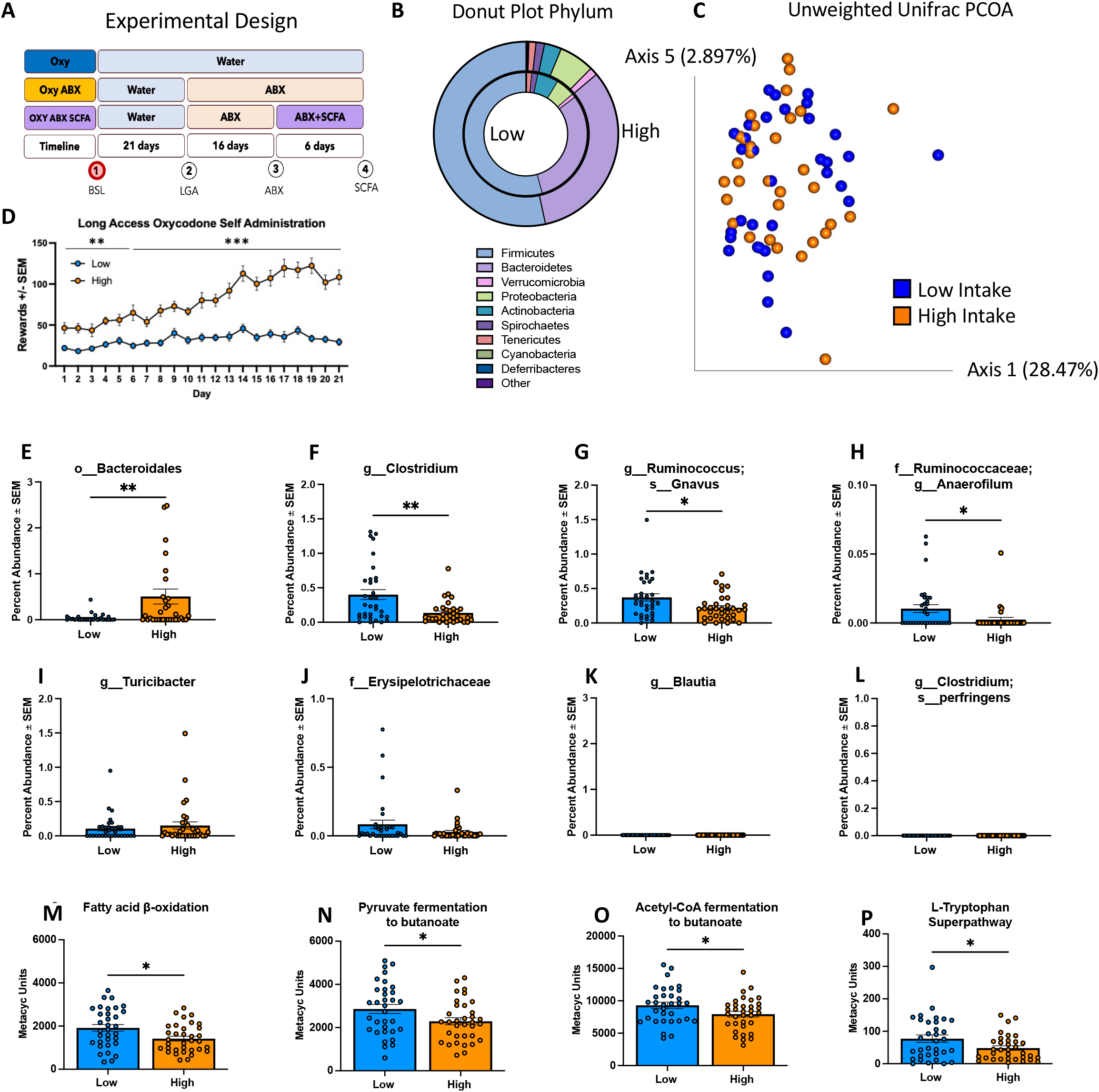
High intake individuals have distinct microbiome and metabolic profiles before drug exposure. Quantification of relevant microbial constituents and pathways. A) Chart depicts the experimental design. Chart is highlighted at BSL, indicating that sampling was taken prior drug exposure. A) Experimental design – Animals were split into three groups, Oxy, Oxy+ABX (ABX), and Oxy+ABX+SCFA (SCFA). All animals received only normal drinking water for the first 21 days of 12h long-access self-administration. In the second phase (16 days), the ABX and ABX+SCFA groups received non-absorbable antibiotics in their drinking water. In the third phase (6 days), the ABX group continues to receive non-absorbable antibiotics, the SCFA group receives non-absorbable antibiotics and a cocktail of short chain fatty acids in their drinking water. The water group receives normal drinking water throughout the duration of the experiment. Feces were harvested at four time-points – 1) BSL (Pre-drug exposure), 2) LgA (Post-acquisition long access self-administration), 3) ABX (Post-ABX exposure), 4) SCFA (Post SCFA-exposure. B) Donut plot at the phylum level of BSL (pre-drug exposure) microbes. C) Unweighted Unifrac PCOA. D) Graph shows oxycodone intake during 12h of long-access self-administration. Animals were separated into high and low intake groups by a median split of the average of the last three days of intake. The high-intake group exhibited a significant increase in oxycodone infusions on Day 1-7 (*p* < 0.0261), and Day 8-21 (p < *0*.001). E-L) Microbial constituents that are altered over the duration of the experiment. The level is proceeding the name – ranging from order to species level. 16S RNA analysis was performed at the phylum, order, and genus level. Unpaired two-tailed students *t*-tests revealed no significant differences at the phylum level, but there are significant increases in *Bacteriodales* (*p* < 0.006, t = 2.836) (Figure 1E), *Ruminoccocus Anaerofilum* (p < 0.018) (Figure 1H), and the species *Ruminococcus Gnavus* (p < 0.015) (Figure 1F). The genus *Clostridium* is significantly lower in high intake individuals (*p* < 0.001, t = 3.425) (Figure 1F). *Turcibacter, Erysipelotrichaceae, Blautia*, and *Clostridium perfringens* are not significantly different between high and low intake individuals at baseline. M-P) Several metabolic pathways that were later revealed to be significant in animals that were sensitive to SCFA administration after antibiotic depletion were also altered prior to drug intake. Unpaired t-tests with Welch’s correction were used to assess the pathways displayed in M-P. High-intake individuals exhibited decreases in Fatty acid β-oxidation (p < 0.013), Pyruvate fermentation to butanoate (p = 0.036), acetyl-CoA fermentation to butanoate (p = 0.042), and Superpathway of Tryptophan Biosynthesis (p = 0.041). *p<0.05, **p<0.01, ***p<0.001. Abbreviations: ABX-Antibiotics, BSL-Baseline pre-drug exposure, LgA – Long Access Self Administration, SCFA-Short Chain Fatty Acids.

### Qiime2 and PICRUSt2

After sequencing, sequences were demultiplexed, and processed using the Quantitative Insights into Microbial Ecology (QIIME2) software package (Bolyen et al., 2019). The DADA2 plugin in QIIME2 (q2-dada2) was used to generate Amplicon Sequence Variants (ASVs). Principal coordinate analysis (PCoA) of the unweighted UniFrac distances was used to assess the variation between samples (beta diversity) (Lozupone et al., 2011). Beta diversity was further visualized via PCoA plots – weighted and unweighted unifrac distances were generated. Taxonomy assignment and rarefaction were performed using QIITA and Qiime2 with 15,000 reads per sample (Gonzalez et al., 2018). The alpha diversity was measured using both the Shannon diversity index and Chao1 index and compared between baseline and treatment conditions. The PICRUSt2 pipeline was used with ASVs to infer predicted sample pathway abundances (Douglas et al., 2020).

### Statistics

The data are expressed as the mean ± SEM. Parametric data with two different groups was assessed with a student’s t-test with Welch’s correction and Bonferroni post-hoc, non-parametric data with two groups was assessed with a Mann-Whitney test. Repeated measures such as rewards were analyzed using a mixed-model ANOVA, with group and time as within-subjects factor. Statistics on behavioral outcomes and corresponding graphs were generated using PRISM 9 Graphpad (San Diego, California, USA). Diversity plots and relative abundance stacked bar charts were generated via QIIME. Differences in distinct clustering in PCA plots was assessed via PERMANOVA method using vegan R-package through QIIME2. Values of *p* < 0.05 were considered statistically significant.

## Results

### High intake individuals exhibit distinct microbiome profiles prior to drug exposure and exhibit disruption of metabolic profiles associated with SCFA and Tryptophan production

The experimental design is depicted in **Figure 1A**. Male and Female HS rats (n = 68 [34 males, 34 females]) were allowed 12 hours of long access (LgA) self-administration of oxycodone for 21 days to acquire and stabilize oxycodone intake. The final three days of LgA were averaged and the median split was taken to determine if individuals are considered part of the “High” intake group or “Low” intake group. A two-way RM-ANOVA revealed a significant interaction (time x group) (*F*_20,660_ = 10.31, *p* < 0.001). The Bonferroni *post-hoc* test revealed increased intake in the High intake group compared to the Low intake group at days 1-3 (*p* < 0.0261), days 4-5 (*p* < 0.05), days 6-7 (*p* < 0.003) and days 8-21 (*p* < 0.001). High intake individuals continued to escalate intake over the 21-day period, whereas low intake individuals stabilized around day 7-8 and did not escalate intake (**Figure 1D**).

The donut plot (**Figure 1C)** displays the taxonomic relative abundance at the phylum level at BSL (pre-drug exposure). 16S RNA analysis was performed to determine the taxonomic relative abundances of the resident microbiota. Unpaired two-tailed students t-tests revealed no significant differences at the phylum level prior to drug exposure. Despite no differences at the phylum level, distinct profiles of individual taxa emerged from the two groups. High intake individuals exhibited an increased relative abundance of *Bacteriodales* (*p* < 0.006, t = 2.836) (**Figure 1E**), *Ruminoccocus Anaerofilum* (*p* < 0.018, t = 1.732) (**Figure 1H**), and *Ruminococcus Gnavus* (*p* < 0.015, t = 0.4814) compared to the Low intake group (**Figure 1G**). High intake individuals exhibited a lower relative abundance in the genus *Clostridium* compared to low intake individuals (*p* < 0.001, t = 3.425) (**Figure 1F**). The genus *Turcibacter*, the family *Erysipelotrichaceae*, the genus *Blautia*, and *Clostridium perfringens* are not significantly different between High and Low intake individuals at the pre-drug exposure collection timepoint (baseline) (**Figures 1M-P**); however, differences are observed after oxycodone access **(Figure 2 H-K**) and are tracked throughout the treatment epochs of the experiment.

**Figure 2.**
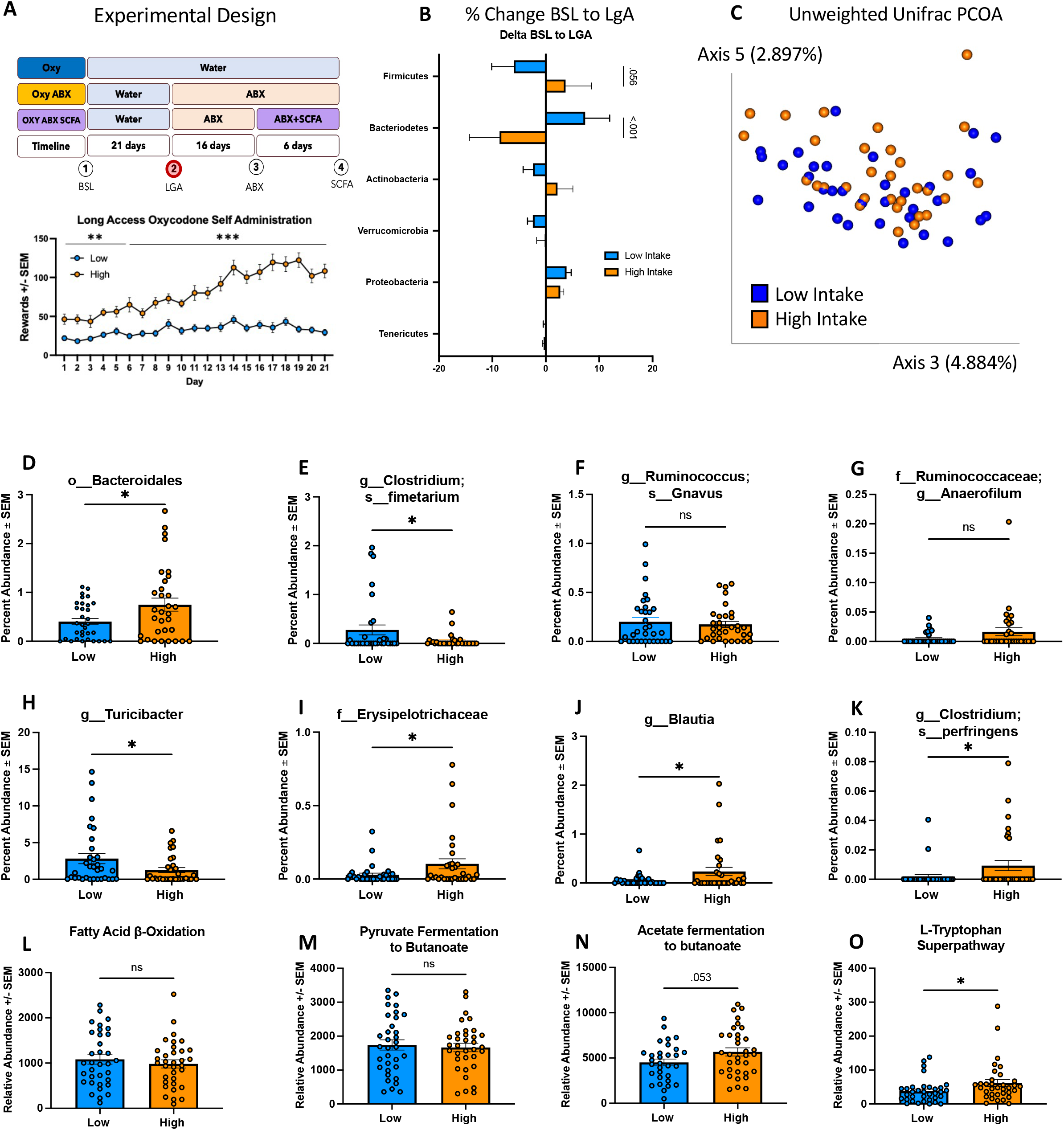
Oxycodone Exposure decreases Bacteroides and increases Firmicutes, Actinobacteria and Proteobacteria in High intake individuals. A) Chart depicts the experimental design. Chart is highlighted at LgA, indicating that sampling was taken after completion of Day 21 of LgA. Graph shows Oxycodone intake during 12h of long-access self-administration. Animals were separated into High and Low intake groups by a median split of the average of the last three days of intake. The high-intake group exhibited a significant increase in oxycodone infusions on Day 1-7; p < 0.0261, and Day 8-21; p < 0.001. B) Bar plot displays the percent change in key microbial populations from BSL to LgA in low and high intake individuals. A two-way RM ANOVA revealed a significant interaction (group x time) (*p* < 0.001). The Bonferroni post-hoc test revealed high intake individuals exhibited an increase in Firmicutes that trends towards significance (p = 0.056). High intake individuals exhibited a significant decrease in *Bacteriodetes* (p < 0.001). No significant change occurred in *Actinobnacteria, Verrucomicrobia, Proteobacteria*, and *Tenericutes* between High and Low intake individuals from BSL to LgA. C) Unweighted Unifrac displaying distribution of high and low intake individuals D-K) Microbial constituents that are altered over the duration of the experiment. The level is proceeding the name – ranging from order to species level. High intake individuals exhibited significant increases in *Bacteriodales* (p = 0.022), *Erysipelotrichaceae* (*p* = 0.040), *Blautia* (*p* = 0.039), and *Clostridium perfringens* (p = 0.047). High intake individuals exhibited significant decreases in *Clostridium fimetarium* (p = 0.035) and *Turicibacter* (p = 0.047). There were no significant changes in *Ruminococcus Gnavus* or *Ruminococcaceae Anaerofilum*. L-O) High-intake individuals exhibited a significant increase in L-Tryptophan Superpathway (*p* = 0.011), and an increase that trended towards significance in Acetate fermentation to butanoate (*p* = 0.053). No significant differences were observed in Fatty acid β-oxidation or Pyruvate fermentation to butanoate (*p* > 0.05). *p<0.05, **p<0.01, ***p<0.001. Abbreviations: ABX-Antibiotics, BSL-Baseline pre-drug exposure, LgA – Long Access Self Administration, SCFA-Short Chain Fatty Acids.

PICRUSt2 analysis of the 16S rRNA sequencing revealed several MetaCyc pathways that are related to SCFA and tryptophan metabolism. A student’s t-test was used to assess the pathways displayed in **Figures 1M-P**. High-intake individuals exhibited less expression in fatty acid β-oxidation (*p* < 0.013, t = 2.571) (**Figure 1M**), Pyruvate fermentation to butanoate (*p* = 0.036, t = 2.145) (**Figure 1N**), Acetyl-CoA fermentation to butanoate (*p* = 0.042, t = 2.079) (**Figure 1O**), and the Superpathway of tryptophan biosynthesis (*p* = 0.041, t = 2.079) (**Figure 1P**). Fatty acid catabolism (fatty acid β-oxidation) and fermentation of molecules to produce butanoate (butyrate, a SCFA) are essential steps in the secretion of SCFAs into the circulating blood stream.

### Oxycodone exposure decreases Bacteroides and increases Firmicutes, Actinobacteria and Proteobacteria in the high intake group

**Figure 2A** represents the sampling timepoint after 21 days long access self-administration (2). The bar plot in **Figure 2B** displays the percent change in key microbial populations from BSL to LgA in high and low intake groups. A two-way RM ANOVA revealed a significant interaction (phylum x group) (*F*_5,198_ = 5.524, *p* < 0.001). The Bonferroni *post-hoc* test revealed the high intake group exhibits an increase in *Firmicutes* that trends towards significance (*p* = 0.056) compared to the low intake group. The High intake group also exhibits a significant decrease in *Bacteriodetes* (*p* < 0.001) compared to the Low intake group. No significant change occurred in *Actinobacteria, Verrucomicrobia, Proteobacteria*, and *Tenericutes*. The PCOA of Unweighted Unifrac distance is useful for examining differences in low-abundance features - **Figure 2C** depicts the distribution of high and low intake individuals.

**Figures 2D-K** plot microbial constituents that are altered over the duration of the experiment. Unpaired two-tailed students t-tests revealed high intake individuals experienced significant increases in the order *Bacteriodales* (*p* = 0.022, *t* = 2.355) (**Figure 2D**), the family *Erysipelotrichaceae* (*p* = 0.040, *t* = 2.099) (**Figure 2I**), the genus *Blautia* (*p* = 0.039, *t* = 2.112) (**Figure 2J**), as well as *Clostridium perfringens* (*p* = 0.047, *t* = 2.023) (**Figure 2K**). High intake individuals exhibit significant decreases in *Clostridium fimetarium* (*p* = 0.035, *t* = 2.151) (**Figure 2E**) and the genus *Turicibacter* (*p* = 0.047, *t* = 2.023) (**Figure 2H**). There were no significant changes (*p* > 0.05) in *Ruminococcus Gnavus* (**Figure 2F**) or *Ruminococcaceae Anaerofilum* (**Figure 2G**).

For Metacyc pathways involved in SCFA and Tryptophan production, the High-intake group exhibit a significant increase in the Superpathway of tryptophan biosynthesis (*p* = 0.019, Mann-Whitney U = 387.5) (**Figure 2O**), and an increase that trended towards significance in Acetyl-CoA fermentation to butanoate (*p* = 0.053, t = 1.976) (**Figure 2N**). No significant differences were observed in Fatty acid β-oxidation (**Figure 2L**) or Pyruvate fermentation to butanoate (**Figure 2M**) (*p* > 0.05).

### Antibiotic depletion increases Verrucomicrobia and Proteobacteria and decreases Bacteroidetes and Firmicutes at the phylum level. Fatty acid β-oxidation and the Tryptophan Superpathway is increased after abx depletion while SCFA pathways are decreased

Fecal samples were taken after antibiotic treatment (Fig 3A) timepoint 3 (red dot). A 2-way mixed effects ANOVA revealed animals that were either put into the water or abx groups were comparable in terms of rewards during self-administration on days 1-21 (p > 0.05) The bar plot in **Figure 3B** displays the percent change in key microbial populations from LgA to ABX in the ABX and Water groups. A mixed-effects ANOVA revealed no significant interaction (time x group) F_(20, 397)_ = 1.178, p = 0.270). There was a significant effect of time (F _(20, 880)_ = 16.99, *p* < 0.001), but no effect of group (F _(1, 44)_ = 0.0002838, *p* = 0.987). The ABX group exhibit significant increases in the phyla *Verrucomicrobia* (*p* = 0.002) and *Proteobacteria* (*p* = 0.008). ABX individuals exhibit a significant decrease in *Firmicutes* (*p* = 0.011) and *Bacteriodetes* (*p* < 0.001). No significant differences occurred in *Fusobacteria, Actinobacteria*, and *Spirochaetes*. The PCOA of Unweighted Unifrac reveals significant separation between Water and ABX groups after ABX exposure (*p* < 0.05) (**Figure 3C**).

**Figure 3.**
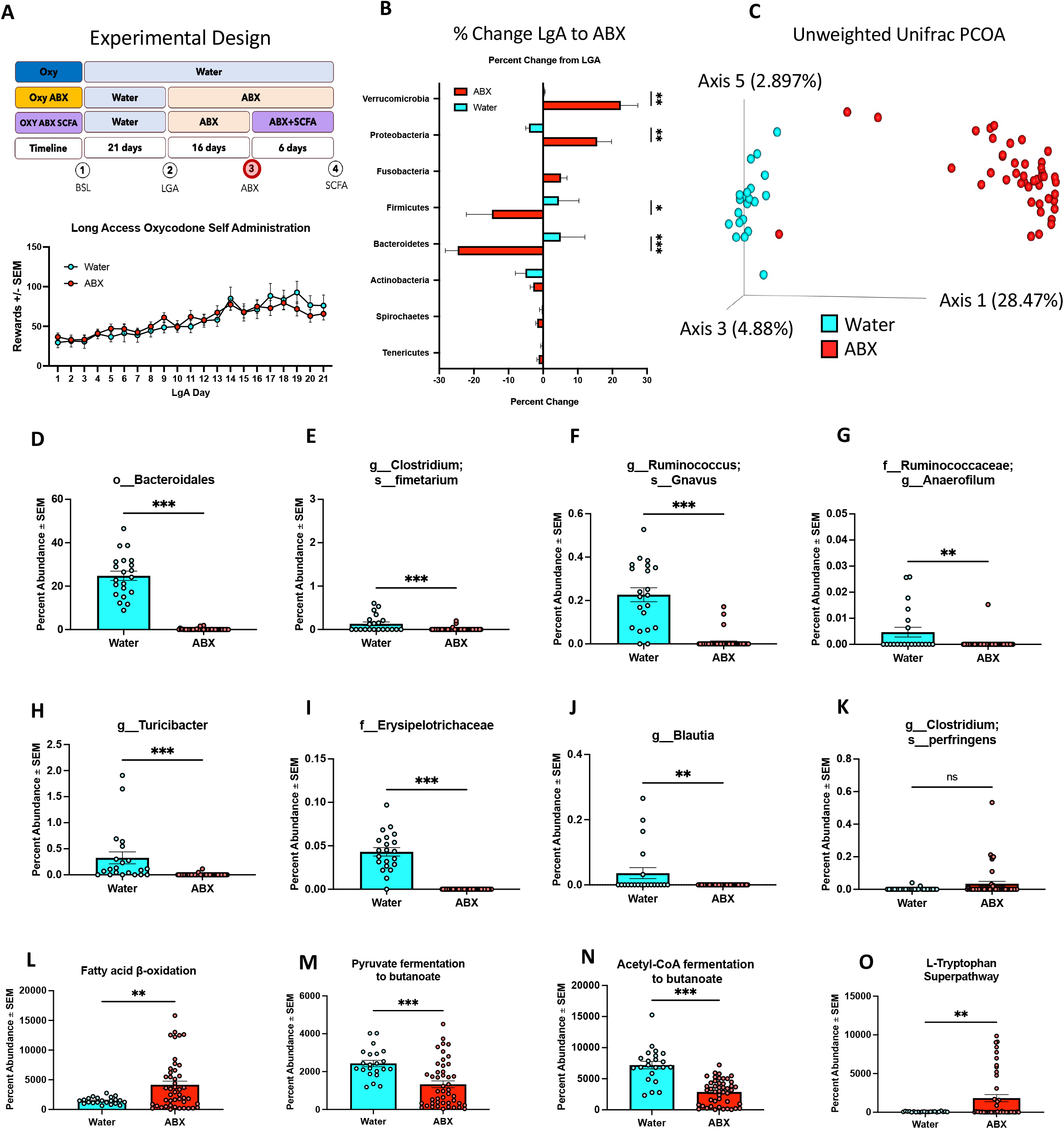
Antibiotic depletion increases Verrucomicrobia, proteobacteria, and decreases Bacteroidetes and firmicutes. A) Chart depicts the experimental design. Chart is highlighted at ABX, indicating that sampling was taken after ABX exposure. Bottom graph displays Oxycodone intake during 12h of long-access self-administration. Animals were separated into Water and ABX groups. A 2-way mixed effects ANOVA revealed animals before ABX treatment were comparable to water group. B) Bar plot displays the percent change in key microbial populations from LgA to ABX in ABX and Water groups. A mixed-effects analysis revealed a significant interaction (Time x Group) (p < 0.001). ABX individuals exhibited significant increases in *Verrucomicrobia* (p = 0.002) and *Proteobacteria* (p = 0.008). ABX individuals exhibited a significant decrease in *Firmicutes* (p = 0.011) and *Bacteroidetes* (p < 0.001). No significant change occurred in Fusobacteria, *Actinobacteria*, and *Spirochaetes*. C) Unweighted Unifrac shows separation of water and ABX group D-K) ABX group exhibited decreases in key microbial constituents. ABX group had significant decreases in *Bacteriodales* (p < 0.001), *Clostridium fimetarium* (p < 0.001), *Ruminococcus gnavus* (p < 0.001), *Ruminococcus Anaerofilum* (p = 0.002), *Turicibacter* (p < 0.001), *Erysipelotrichaceae* (p < 0., t = 13.08), and *Blautia* (p < 0.003). No significant difference was observed in *Clostridium perfringens*. L-O) ABX individuals exhibited a significant increase in Fatty acid β-oxidation (p = 0.004) and the L-Tryptophan Superpathway (p = 0.010). ABX individuals exhibited a significant decrease in Pyruvate fermentation to butanoate (p < 0.001) and Acetyl-CoA fermentation to butanoate (p < 0.001). *p<0.05, **p<0.01, ***p<0.001. Abbreviations: ABX-Antibiotics, BSL-Baseline pre-drug exposure, LgA – Long Access Self Administration, SCFA-Short Chain Fatty Acids.

The ABX group exhibit decreases in the key microbial constituents that have been demonstrated to be altered throughout the experiment. An unpaired t-test revealed the ABX group had significant decreases in *Bacteriodales* (*p* < 0.001, *t* = 16.80) (**Figure 3D**), *Clostridium fimetarium* (*p* < 0.001, *t* = 4.151) (**Figure 3E**), *Ruminococcus gnavus* (*p* < 0.001, *t* = 9.386) (**Figure 3F**), *Ruminococcus Anaerofilum* (*p* = 0.002, *t* = 3.175) (**Figure 3G**), *Turicibacter* (*p* < 0.001, *t* = 4.120) (**Figure 3H**), *Erysipelotrichaceae* (*p* < 0.001, *t* = 13.08) (**Figure 3I**), and *Blautia* (*p* < 0.003, *t* = 3.108) (**Figure 3J**). No significant difference was observed in *Clostridium perfringens* following antibiotic depletion (**Figure 3K**).

PICRUSt2 MetaCyc pathway analysis demonstrates significant differences between the Water group and the ABX group. Contrary to the high intake group at baseline, the ABX group exhibit a significant increase in Fatty acid β-oxidation (*p* = 0.004, *t* = 2.944) (**Figure 3L**) and L-Tryptophan Superpathway (*p* = 0.010, *t* = 2.654) (**Figure 3O**). Similar to the High intake group at baseline, The ABX group exhibit a significant decrease in Pyruvate fermentation to butanoate (*p* < 0.001, *t* = 3.910) (**Figure 3M**) and Acetyl-CoA fermentation to butanoate (*p* < 0.001, *t* = 7.270) (**Figure 3N**).

### Antibiotic Depletion Exacerbates Oxycodone Escalation. Responders exhibit distinct microbial profiles

**Figure 4A** depicts the fecal sampling timepoint in the experimental design (After ABX treatment). **Figure 4D** plots the percent change of the rewards taken by the Responder and Non-Responder groups during the ABX treatment were calculated by a change from the average of the last three days of LgA (Day 19, 20, 21) to each day of ABX treatment. Animals were separated into Responder and Non-Responder groups by comparing the last three days of LgA prior to ABX treatment (Day 19, 20, 21), to the last three days of ABX treatment. Animals were considered Responders if there was an increase from the first timepoint to the second timepoint. A 2-way REML ANOVA did not reveal an interaction (time x group) F_(3.856, 28.02)_ = 1.788 (p = 0.161) or (time) F_(3.885, 136.0)_ = 1.425, but did show a significant increase of (group) F_(1.000, 35.00)_ = 4.760. The Bonferroni *post-hoc* significantly higher values in Responders at day 3 (p = 0.001), day 4 (p = 0.015), day 5 (*p* = 0.008), day 6 (*p* = 0.0023), day 7 (*p* = 0.006), day 8 (*p* = 0.003), day 9 (*p* < 0.001), day 10 (*p* = 0.020), day 11 (*p* = 0.121), day 12 (*p* = 0.014), day 13 (*p* = 0.006), day 14 (*p* = 0.003), day 15 (*p* = 0.002), day 16 (*p* = 0.003).

**Figure 4.**
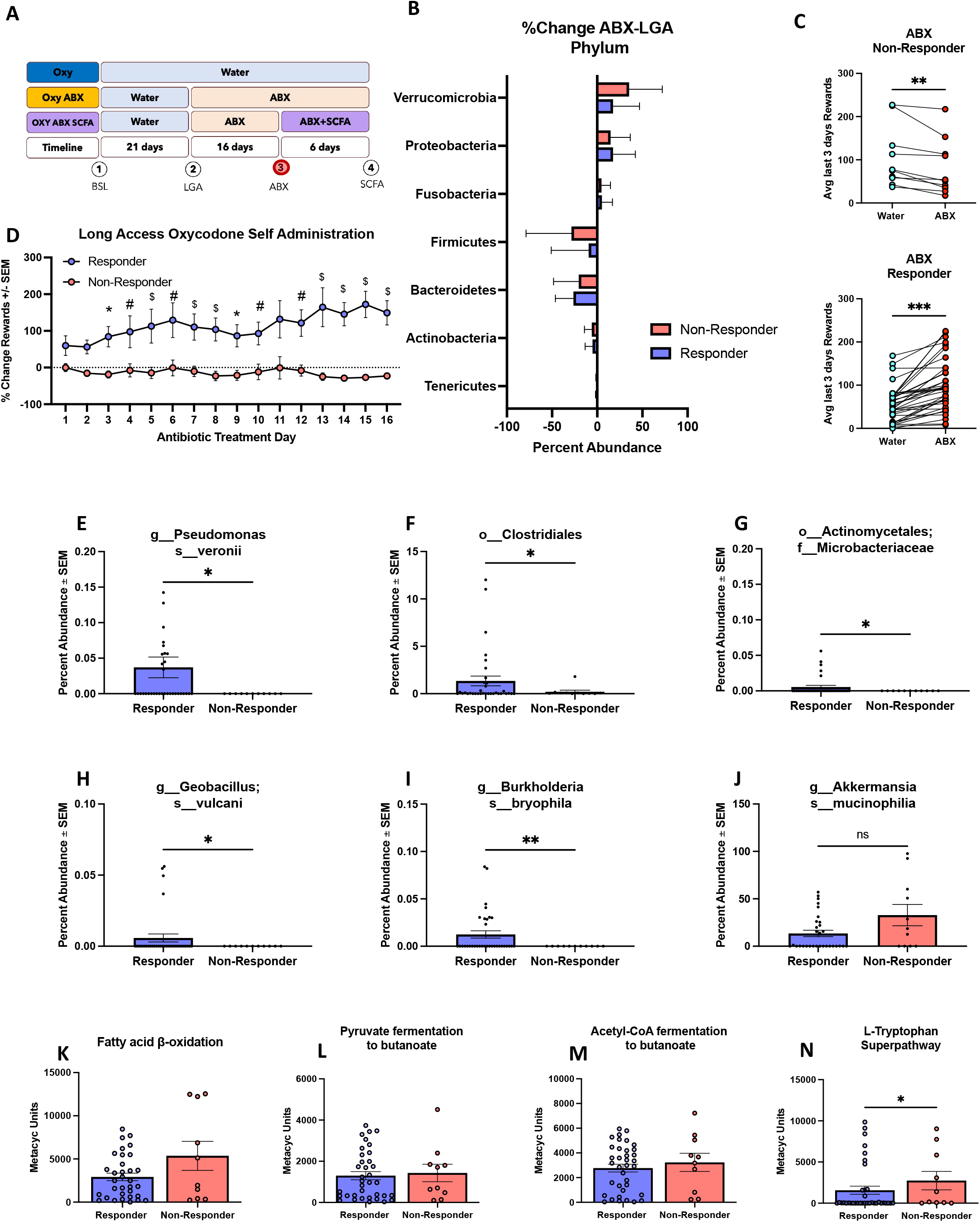
Antibiotic Depletion Exacerbates Oxycodone Intake. Responders exhibit distinct microbial profiles. A) Experimental design B) Bar plot displays the change in key microbial phylum between ABX and LgA in Responder and Non-Responder groups. No significant change was observed in *Verrucomicrobia, Proteobacteria, Fusobacteria, Firmicutes, Bacteroidetes, Actinobacteria, Spirochaetes*, and *Tenericutes*. C) Figure 4C plots average of rewards taken during the last three days of long access (Day 19, 20, 21) prior to antibiotic treatment, and the last three days of ABX treatment (Day 13, 14, 15). Non-Responders showed a significant decrease in rewards between Water and ABX timepoint (*p* =0.006). D) Figure 4D plots the percent change of the rewards taken by the Responder and Non-Responder groups during the ABX epoc of long-access self-administration calculated by a change from the average of the last three days of long access self-administration to each day of ABX treatment. Animals were separated into Responder and Non-Responder groups by comparing the last three days of long-access prior to ABX treatment, to the last three days of ABX treatment. Animals were considered Responders if there was an increase from the first timepoint to the second timepoint. A 2-way mixed effects ANOVA revealed animals in Responder groups experienced a significant increase in intake (*p* < 0.05). E-J) Microbial constituents that are altered over the duration of the experiment. The level is proceeding the name – ranging from order to species level. A t-test with Welch’s correction was used to assess graphs E-J. Responder individuals exhibited significant increases in *Clostridiales* (p = 0.040, t = 2.125), the family *Actinomycetales Microbacteriaceae* (p = 0.031), and the species Pseudomonas veronii (p = 0.016, t = 2.532), *Geobacillus vulcani* (p = 0.046, t =2.070), and *Burkholderia bryophila* (p = 0.003, t = 3.185). There were no significant changes in *Akkermansia mucinophilia*. K-N) L-Tryptophan Superpathway was elevated in the Non-Responder group (*p* = 0.041). There were no significant differences in Fatty acid β-oxidation, Pyruvate fermentation to butanoate, and Acetyl-CoA fermentation to butanoate (Figures 4K-M) (*p* > 0.05). *p<0.05, **p<0.01, ***p<0.001. Abbreviations: ABX-Antibiotics, BSL-Baseline pre-drug exposure, LgA – Long Access Self Administration, SCFA-Short Chain Fatty Acids.

The bar plot in **Figure 4B** displays the change in key microbial phylum in the Responder and Non-Responder groups comparing before and after antibiotic treatment (ABX – LgA percent abundance). No significant change was observed in *Verrucomicrobia, Proteobacteria, Fusobacteria, Firmicutes, Bacteriodetes, Actinobacteria, Spirochaetes*, and *Tenericutes*. Both Responder and Non-responder groups were treated with ABX to deplete the gut microbiota.

**Figure 4C** plots average of rewards taken during the last three days of LgA (Day 19, 20, 21) prior to antibiotic treatment, and the last three days of ABX treatment (Day 13, 14, 15). Non-Responders showed a significant decrease in rewards between Water and ABX timepoint (*p* = 0.006, *t* = 3.564). ABX Responders showed a significant increase in rewards between the Water and ABX timepoint (*p* < 0.001, *t* = 6.861).

**Figures 4E-J** display the microbial constituents that are significantly different between the Responder and Non-Responder groups. The Responder group exhibited significant increases in the order *Clostridiales* (*p* = 0.040, *t* = 2.125) (**Figure 4F**), *Actinomycetales Microbacteriaceae* (*p* = 0.031, *t* = 2.250) (**Figure 4G**), and *Pseudomonas veronii* (*p* = 0.016, *t* = 2.532) (**Figure 4E**), *Geobacillus vulcani* (*p* = 0.046, *t* =2.070) (**Figure 4H**), and *Burkholderia bryophila* (*p* = 0.003, *t* = 3.185) (**Figure 4I**). There were no significant changes in the species *Akkermansia mucinophilia* (*p > 0*.*05)* (**Figure 4J**).

**Figures 4K-N** display several MetaCyc pathways that were later revealed to be significant in animals that following antibiotic depletion. The Tryptophan Superpathway was elevated in the Non-Responder group (*p* = 0.041, *Mann-Whitney U* = 104) (**Figure 4N**). There were no significant differences in Fatty acid β-oxidation, Pyruvate fermentation to butanoate, and Acetyl-CoA fermentation to butanoate (**Figures 4K-M**) (*p* > 0.05).

### SCFA administration partially rescues ABX induced oxycodone intake escalation

**Figure 5A** depicts the experimental timepoint; however, all sequencing for this timepoint occurred before SCFA treatment and after ABX treatment. The SCFA cocktail was administered between timepoint 3 and 4 for 6 days to animals that received the ABX treatment. **Figure 5B** plots SCFA Non-Responder and Responder groups at the average of the last three days of the ABX timepoint and the last three days of SCFA treatment. Animals were considered Non-Responders to the SCFA cocktail if they did not exhibit a decrease in an average of the rewards in the last three days of the SCFA treatment compared to the last three days of the ABX treatment. Animals were considered Responders to the SCFA cocktail if they exhibited a decrease in the average of rewards in the last three days the SCFA treatment compared to the last three days of the ABX treatment. SCFA Non-Responders showed an increase between SCFA and ABX timepoint following treatment that trended towards significance (*p* = 0.057, *t* = 2.279). SCFA Responders showed significant decrease following SCFA treatment between SCFA and ABX timepoint (*p* = 0.014, *t* = 2.873). **Figures 5C-J** display the MetaCyc pathways that exhibit significant differences between the group that responded to SCFA treatment (Responder), and the group that did not (Non-Responder). The SCFA Responder group had significant increases in Pyruvate fermentation to acetone (*p* = 0.018, *t* = 2.611) (**Figure 5C**), Adenosine nucleotide degradation II (*p* = 0.033, *t* = 2.308) (**Figure 5D**), Formaldehyde assimilation II (*p* = 0.049, *t* = 2.105) (**Figure 5E**), L-leucine degradation I (*p* = 0.049, *t* = 2.135) (**Figure 5G**), L-arginine degradation VI (*p* = 0.050, *t* = 2.098) (**Figure 5H**), Allantoin degradation to glyoxylate III (*p* = 0.050, *t* = 2.152) (**Figure 5I**), and a decrease in Nucleoside and nucleotide degradation that trended towards significance (*p* = 0.061, *t* = 1.998) (**Figure 5J**). The SCFA Responder group experienced a significant decrease in Sucrose degradation III compared to the Non-Responder group (*p* = 0.037, *t* = 2.487) (**Figure 5F**). **Figures 5K-N** display the MetaCyc pathways that were differentially abundant in the High intake group prior to drug exposure. Fatty acid β-oxidation appeared reduced in Responders but was not significant (p = 0.356, t = 0.9637) **(Figure 5K)**, Pyruvate fermentation to butanoate was not significantly different between the two groups (p = 0.251, t = 1.187) **(Figure 5L)**, Acetyl-CoA fermentation to butanoate was not significantly different between the two groups (p = 0.165, t = 1.446) (**Figure 5M)**, and L-Tryptophan Superpathway was elevated in the Responder group compared to the Non-Responder group but did not achieve significance (p = 0.115, t = 1.652) **(Figure 5N)**.

**Figure 5.**
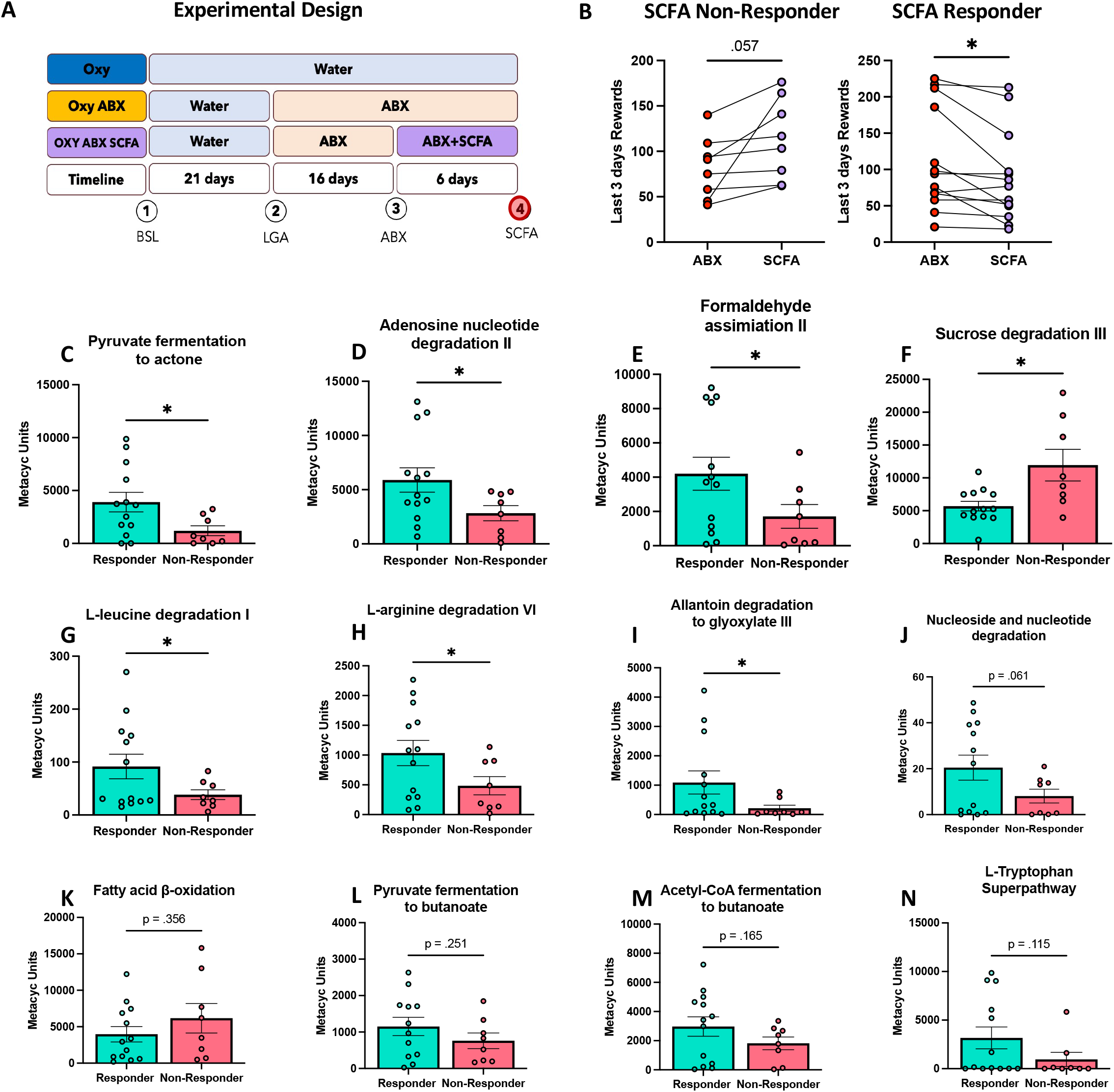
SCFA administration partially rescues ABX induced escalation. A) Experimental Design. B) Figure 5B plots SCFA Non-Responder and Responder groups at the average of the last three days of the ABX timepoint and the last three days of SCFA treatment. Animals were considered Non-Responders to the SCFA cocktail if they did not exhibit a decrease in an average of the rewards in the last three days of the SCFA epoc compared to the last three days of the ABX epoc. Animals were considered Responders to the SCFA cocktail if they exhibited a decrease in the average of rewards in the last three days the SCFA epoc compared to the last three days of the ABX epoc. SCFA Non-Responders showed an increase between SCFA and ABX timepoint following treatment that trended towards significance (*p* = 0.057). SCFA Responders showed significant decrease following SCFA treatment between SCFA and ABX timepoint (*p* = 0.014). C-E) The SCFA Responder group had significant increases in Pyruvate fermentation to acetone (*p* = 0.018, *t* = 2.611) (Figure 5C), Adenosine nucleotide degradation II (*p* = 0.033), Formaldehyde assimilation II (*p* = 0.049), L-leucine degradation I (*p* = 0.049), L-arginine degradation VI (*p* = 0.050) (Figure 5H), Allantoin degradation to glyoxylate III (*p* = 0.050), and a decrease in Nucleoside and nucleotide degradation that trended towards significance (*p* = 0.061). The SCFA Responder group experienced a significant decrease in Sucrose degradation III compared to the Non-Responder group (*p* = 0.037). Fatty acid β-oxidation was elevated in Non-Responders and trended towards significance (p = 0.356, t = 0.9637), Pyruvate fermentation to butanoate was decreased in Non-Responders and trended towards significance (p = 0.251), Acetyl-CoA fermentation to butanoate was not significant (p = 0.165), and L-Tryptophan Superpathway was elevated but not significant (p = 0.115).

## Discussion

This study identifies pre-existing microbiome and metabolic vulnerabilities to escalation of oxycodone self-administration, demonstrates that escalation of oxycodone self-administration dysregulates the microbiome and metabolic landscape, and identifies a causal role of short-chain fatty acids in addiction-like behaviors. We found distinct microbial and metabolic functions prior to exposure to opioids in individuals that ultimately escalated oxycodone intake (High intake group) when given long access (12 h/day) to oxycodone self-administration. The results also demonstrates that escalation of oxycodone self-administration dysregulates the microbiome and metabolic landscape and identifies a causal protective role of short-chain fatty acids in against addiction-like behaviors.

Characterization of the microbiome in the High intake group prior to drug intake was to identify pre-existing vulnerabilities in microbiome contributions and constituents that may lead to escalation of intake and to determine if the same profiles were present after antibiotic depletion of the microbiome. The unweighted unifrac PCOA demonstrated slight separation of the groups; however, since there is a median split dictated by behavior, there are individuals that are bound to be in separate groups with similar profiles contributing to the overlap. While at the phylum level, there appears to be no difference between the High and Low intake groups, some individual differences in taxa relative abundance suggests that bacterial communities like *Bacteriodales* which are known to be a primary producer of the SCFA propionate within the gut microbiome (Reichardt et al., 2014) highlighting the importance of thorough taxonomic profiling. A reduced relative abundance of the genus *Clostridium* (another SCFA producer) suggests potential dysregulation of SCFA production in High Intake group prior to drug exposure. One species (Gnavus) and one genus (Anaerofilum) of Ruminococcus exhibited decreased relative abundance in the High intake group - increased levels of *Ruminococcus gnavus*, a mucin degrader from glucose and malto-oligosaccharides released by the keystone species *Ruminococcus Bromii*, which is important for degrading resistant starches in the colon (Crost et al., 2018). In co-culture experiments an increase in *Ruminococcus gnavus*, causes *Ruminococcus Bromii* to upregulate tryptophan biosynthesis genes and down regulate vitamin B-12 dependent methionine biosynthesis, suggesting usage of SCFA and increased levels of alternative tryptophan metabolism in individuals with elevated *Ruminococcus gnavus* (Crost et al., 2018). Indeed, the Low intake group has increased relative abundance of the tryptophan super pathway in comparison to the High intake group. Additionally, high intake of sugar is linked it lower relative abundances of *Anaerofilum* in obese patients (Koo et al., 2019) and similar increases in preference for sweet taste is associated with glycemic dysregulation in chronic opioid users (Mysels and Sullivan, 2010). Perhaps the beginnings of craving preference and tryptophan availability are set prior to opioid exposure and contribute to later escalation.

The MetaCyC pathway analysis exhibited decreased abundance of Fatty acid b-oxidation, pyruvate fermentation to butanoate, Acetyl CoA fermentation to butanoate, and L-Tryptophan superpathway in the High intake group. An abundance of SCFAs has been demonstrated to stimulate fatty acid oxidation through histone-deacetylase gene regulation, activating UCP2-AMPK signaling – in fact increased butyrate availability increases lipid metabolism as a positive feedback loop and are tightly associated with signaling pathways in inflammation, glucose and lipid metabolism (He et al., 2020). Perhaps a reduction of fermentation of pyruvate and Acetyl-CoA to butanoate (butyrate) in high intake individuals reduces available SCFAs to properly regulate fatty acid oxidation and downstream metabolic regulation signaling involved in energy homeostasis and craving. Finally, a reduction in tryptophan synthesis has been observed in predicting OUD diagnosis in high/chronic opioid users (Ghanbari et al., 2021), and acute tryptophan depletion blocks morphine analgesia (Abbott et al., 1992) while loading of tryptophan has been demonstrated to reduce tolerance by potentially increasing serotonergic pathways (Hosobuchi et al., 1980). Taken together, dysregulation of both SCFA signaling and tryptophan availability may be apparent in individuals prior to escalation of opioids via microbiome or metabolic profiles that appear to shift the importance of SCFA availability to tryptophan availability in the antibiotic Responder group.

Long Access opioid exposure had a significant impact on the microbiome with an increase in the phylum *Firmicutes* and a decrease in *Bacteroidetes* in high intake individuals. *Actinobacteria* and *Proteobacteria*, two proinflammatory phyla were also increased in high intake individuals. Similar to the pre-drug exposure timepoint (1), *Bactoeridales* was elevated and *Clostridium fimentarium* was decreased in the High intake group. Neither *Ruminococcus Gnavus* or *Ruminococcus Anaerofilum* were decreased in the High intake group. Several other microbial constituents were differentially abundant - the family *Turicibacter* was decreased in the High intake group – following oxycodone self-administration. Members of *Turicibacter*, including *Turicibacter sanguinis* have been demonstrated to alter intestinal gene pathways involved in steroid and lipid metabolism, and have a functional serotonergic import system that can reduce available serotonin in the periphery (Fung et al., 2019), *Erysipelotrichaceae, Blautia*, and *Clostridium Perfringens* exhibited higher relative abundance in the High Intake group. Increased relative abundance of *Blautia* has been previously correlated with substance use disorders regardless of age in a study that explored substances that included heroin and methamphetamine (Xu et al., 2017). *Clostridium Perfringens* is observed in healthy adults but is most observed as an opportunistic pathogen in long term care faculties and is elevated levels in heroin users, causing gangrene and food poisoning (Wurcel et al., 2015). Perhaps elevated levels are associated with a host that is at risk of opportunistic pathogens, or functions as a biomarker of impending dysbiosis. MetaCyc pathways after LgA are less dysregulated than what was observed prior to drug exposure – there is no significant difference in Fatty acid β-oxidation, or Pyruvate fermentation to butanoate, but Acetyl-CoA fermentation to butanoate trends towards an elevated relative expression in the High intake group (p = 0.053). The Tryptophan Superpathway that was decreased prior to drug exposure is slightly, but significantly, increased after LgA. Opioid exposure has been demonstrated to increase diversity of the microbiome in rodent models (data not shown) and in human populations (Xu et al., 2017). Tryptophan is a crucial precursor to serotonergic pathways and dysregulation of serotonin can lead to toxicity and/or depressive states that can contribute to increased drug-taking behaviors. Furthermore, some opioids are associated with serotonin toxicity – due to serotonin transporter (SERT) inhibition. While oxycodone and fentanyl are not inhibitory in the same way that tramadol, dextromethorphan, methadone, and meperidine might be, reports have still linked oxycodone and fentanyl to serotonin toxicity in the clinic, suggesting SERT-independent effects on the serotonin system – potentially mediated by the microbiome – *in vivo (Baldo, 2018)*. Both SCFA and tryptophan regulation by the microbiome appear to contribute to high intake behaviors starting from pre-drug exposure and continuing through long-access self-administration of oxycodone.

Antibiotics and opioids are regularly prescribed in tandem in the clinic following surgery, as well as common ailments. Our method of antibiotic depletion is known to specifically reduce a large proportion of SCFA producers. We hypothesized based on previous literature on cocaine (Kiraly et al., 2016a) depletion of the microbiome and the majority of SCFA producers would result in escalation of opioid intake. To ensure that groups were balanced for intake animals were evenly distributed to Water and ABX groups by the last three days of LgA intake (Figure 2A). Antibiotic depletion resulted in significant increases in *Verrucomicrobia, Proteobacteria*, and decreases in *Bacteroidetes* and *Firmicutes*. The unweighted unifrac PCOA demonstrates significant separation of the two groups. The water group exhibits similar profiles of taxa observed in the Low group in Figure 1 (E-K), suggesting that depletion of the microbiome mimics the microbiome of High Intake individuals. Similarly, fermentation to butanoate via Pyruvate and Acetyl-CoA pathways are both decreased. Increases in Fatty acid β-oxidation may suggest alternative pathways for lipid scavenging due to limited availability of SCFAs as the microbial ecosystem shifts from SCFA energy sources to the sugars to mucosal carbohydrate’s to generate sialic acid and fucose – not only leading to increased inflammation but also providing a niche for opportunistic pathogens (Huang et al., 2015; Ng et al., 2013). The Tryptophan superpathway is increased in antibiotic depleted animals resembling High intake individuals after LgA oxycodone exposure.

Depletion of the microbiome exacerbates opioid escalation in a sub-population of animals (Responders). Antibiotic depletion results in nearly a 200% increase on average by the final day of the antibiotic treatment phase compared to the last three days of oxycodone intake prior to antibiotic treatment. No specific microbiome differences were observed at the phylum level (Figure 3B) as the group number is smaller and unbalanced, making detection of distinct differences difficult (n = 10 Non-Responders, n = 36 Responders). Rewards for the last three days of LgA prior to ABX treatment are shown in Figure 3C – Non-Responders significantly decrease intake after ABX treatment, while Responders significantly increase intake. Despite a significant decrease in microbes following ABX treatment, there are still several taxa that exhibit differences between the Responder and Non-Responder groups. *Pseudomonas sp, Clostridiales, Actinomycetales Microbacteriaceae, Geobacillus Vulcani*, and *Bulkholderia Bryophyla* were all significantly increased in the Responder group. Anti-inflammatory *Akkermansia Mucinophilia* trended toward a decreased abundance in the Responder group. Our antibiotic cocktail included an antifungal (pimaricin) to reduce the potential for fungal overgrowth after ABX depletion; *Bulkholderia Bryophyla* possesses anti-fungal properties as well – an increase in *Bulkholderia* may provide insight to other components of the microbiota that are not assessed in this study but may play a role in the observed behavioral changes (Vandamme et al., 2007). The MetaCyc pathways observed at each timepoint associated were all non-significant except for the Tryptophan superpathway which was still elevated. Perhaps the initiation of escalation is linked to SCFA availability, but the perpetuation of the increase may also be related to tryptophan availability.

Finally, a subpopulation of the ABX treated animals were supplemented with a SCFA cocktail for 6 days (n = 21, 13 Responders, 8 Non-responders) to determine if SCFA depletion by ABX treatment could be rescued by dietary repletion. Responders significantly decreased opioid intake following SCFA repletion (Figure 5B). A range of MetaCyc pathways involved in amino acid degradation were elevated in the Responder group. One pathway of interest is the Allantoin degradation to glyoxylate pathway which is increased in the Responder group. Glyoxylate is essential for the condensation propionyl-CoA to α-hydroxyglutarate (Rabin et al., 1965) and may represent an alternative SCFA metabolic pathway. High fecal concentrations of acetic acid are highly correlated glyoxylate metabolism and the formation of uric acid (Liu et al., 2020). Morphine administration has been demonstrated to upregulate adenosine deaminase and xanthine oxidase resulting in elevated plasma levels of uric acid (Liu et al., 2007) which can lead to gout and increase pain and inflammation. It is difficult to determine if the shift is related to elevated oxycodone intake or depletion of the microbiome resulting in alternative SCFA metabolism, yet both pathways resulted in increased opioid intake that was ameliorated by SCFA treatment. The microbiome is a complex, metabolically active ecosystem that is capable of shifting key metabolic pathways involved in the escalation of drug taking behaviors.

To evaluate the effects of the gut microbiome on addiction-like behaviors, the composition and function of the gut microbiome must be characterized over the duration of conversion from non-dependent to dependent. While there is no single example of a healthy microbiome, those who cluster into the “healthy” category tend to have a large proportion of phyla that secrete short-chain fatty acids (SCFA), metabolic products of anaerobic fermentation of dietary fiber (De Filippis et al., 2016). Butyrate, acetate, and propionate comprise the majority of SCFAs that are secreted by the microbiome (Kimura et al., 2011; Louis and Flint, 2017; van de Wouw et al., 2018). SCFAs multifaceted signaling molecules. SCFAs act as ligands for g-protein coupled receptors found throughout the body and the brain, histone deacetylase inhibitors which modify gene expression profiles, and influence inflammatory responses by reducing the recruitment and migration of immune cells (Jayasimhan and Marino, 2019; Kimura et al., 2011; Müller et al., 2019; van de Wouw et al., 2018; Fellows et al., 2018; Vinolo et al., 2011). Many studies, both preclinical and clinical have linked alterations of the microbiome to drugs of abuse, including opioids (Xiao et al., 2018; Hofford et al., 2019; Wang et al., 2018), cocaine (Cho and Blaser, 2012; Volpe et al., 2014; Kiraly et al., 2016a), alcohol (Engen et al., 2015; Leclercq et al., 2017; Leclercq et al., 2020) and psychostimulants (Simpson et al., 2021b; Cook et al., 2019; Yang et al., 2021). Previous studies have demonstrated increases in conditioned place preference (CPP) for cocaine (Kiraly et al., 2016b) and methamphetamine (Ning et al., 2017) was related to circulating SCFA levels and the microbes that produce them. We have demonstrated that antibiotic-induced microbial depletion alters the recruitment of neuronal ensembles that are activated by oxycodone at both intoxication and withdrawal states (Simpson et al., 2020a). Furthermore, dysbiosis of the microbiome is associated with comorbid psychiatric disorders such as anxiety, depression, and stress responsivity indicating a potential for the microbiome to contribute to addiction liability (Capuco et al., 2020; Madison and Kiecolt-Glaser, 2019).

In summary, several lines of evidence suggested that opioid escalation may be influenced by the gut-brain axis and the present study supports this theory by identifying microbial populations and metabolic profiles that are associated high intake individuals prior to drug exposure. The ability to predict an increased risk to opioid misuse prior to antibiotic/opioid exposures would facilitate personalized medicine efforts including the use of alternative pain management tools, as well as increased monitoring and education about opioid misuse and opioid use disorder. Additionally, this study demonstrate that antibiotic depletion of the microbiome may increase risk of escalating opioid use, and that SCFA administration can ameliorate some side effects of antibiotics depletion. Both SCFA and tryptophan metabolic pathways were disturbed in the High intake and ABX Responder group, but SCFAs dysregulation was less obvious after long access self-administration – potentially because SCFA dysregulation is important in initial escalation of oxycodone intake, and tryptophan metabolism combined with serotonin availability and lipid metabolism may play a role in the perpetuation of the cycle. A limitation of the study is that the prediction of the functional composition of the microbial communities is limited to the database and reference genomes. Further studies to determine functional and circulating metabolites will allow for confirmation of the predicted functions. The use of both sexes in a heterogenous outbred populations of animals that self-administer opioids provide translational relevance to uncover similar individuals in the clinic prior to exposure of opioids and opens a new frontier of uncovering live biotherapeutics, beneficial pro- and prebiotics, and small molecules derived from the microbiome and to treat addiction liability at all stages – including the crucial period before an individual is exposed to the drug.

## Acknowledgements

The UCSD Microbiome Core performed sample extractions and library preparation utilizing protocols and primers published on the Earth Microbiome Project website. This publication includes data generated at the UC San Diego IGM Genomics Center utilizing an Illumina NovaSeq 6000 that was purchased with funding from a National Institutes of Health SIG grant (#S10 OD026929).

## Author Contributions

SS, and OG designed the concept and the experiment. SS, AK, RM, ShS, MB, VE, VS, MI, HJ, TH, LCS, JS performed behavioral and/or computational experiments. SS, AK, EW, GP, RM, NW, HJ, SH, MI, VS, TH, MK, LM, LC, TX, VE, JS, LSW, AAP, GdG, OG contributed to manuscript preparation.

